# Transcriptomic and epigenomic profiling of young and aged spermatogonial stem cells reveals molecular targets regulating differentiation

**DOI:** 10.1101/2020.12.15.422121

**Authors:** Jinyue Liao, Hoi Ching Suen, Alfred Chun Shui Luk, Tin-Lap Lee

**Author notes:** To whom correspondence should be addressed.; Address: Rm 622A, 6/F., Lo Kwee-Seong Integrated Biomedical Sciences Building, The Chinese University of Hong Kong, Shatin, Hong Kong SAR, China.

## Abstract

Spermatogonial stem cells (SSC), the foundation of spermatogenesis and male fertility, possess lifelong self-renewal activity. Aging leads to the decline in stem cell function and increased risk of paternal age-related genetic diseases. In the present study, we performed a comparative genomic analysis of mouse SSC (Oct4-GFP+/KIT-) and differentiating progenitors (Oct4-GFP+/KIT+) isolated from young and aged testes. Our transcriptome data revealed enormous complexity of expressed coding and non-coding RNAs and alternative splicing regulation during SSC differentiation. Further comparison between young and aged SSCs suggested these differentiation programs were affected by aging. We identified aberrant expression of genes associated with meiosis and TGF-β signaling, alteration in alternative splicing regulation and differential expression of specific lncRNAs such as Fendrr. Epigenetic profiling revealed reduced H3K27me3 deposition at numerous pro-differentiation genes during SSC differentiation as well as aberrant H3K27me3 distribution at genes in Wnt and TGF-β signaling upon aging. Finally, aged SSCs exhibited gene body hypomethylation, which is accompanied by an elevated 5hmC level. We believe this in-depth molecular analysis will serve as a reference for future analysis of SSC aging.

## INTRODUCTION

In recent decades, there has been a remarkable delay in the paternal age of reproduction across high-income countries. This phenomenon leads to an increase in infertility rate. One area that contributes to age-related infertility is age-associated decline in spermatogonial stem cells. Spermatogonial stem cells (SSCs) are stem cells in the male germline which form the foundation of male fertility. Animal studies have shown that SSCs undergo an age-associated decline in function. For example, there is a dramatic decrease in the quantity of SSCs in aged mouse testes, which causes testicular regression or atrophy and an increased abundance of Sertoli cell-only tubules (Gosden et al. 1982; Ryu et al. 2006; Wang, Leung, and Sinha-Hikim 1993). In contrast to non-germline stem cells, dysregulation of the SSC genome could cause significant health impacts in the offspring. For instance, animal aging models and human cohort studies revealed an increase in *de novo* mutations in germ cells and mature sperms as males enter advanced age. Such changes may lead to developing complex diseases in the offspring (Sanders et al. 2012). Aberrant epigenetic changes in SSC have also been indicated in transgenerational genetic diseases (Guerrero-Bosagna and Skinner 2014; Skinner 2014). Therefore, studying the role of aging in deterioration of SSC function is crucial in the understanding of the aging-associated fertility decline and the impact of delayed parental age to the health of offspring.

Current data suggest that both stem cell intrinsic defects and alterations in microenvironment are implicated in the age-dependent decline of functional germ cell reserves (Boyle et al. 2007; Ryu et al. 2006; Schmidt et al. 2011). Transcriptome studies indicated altered gene expression in aged SSCs, affecting functional pathways such as DNA damage responses, mitosis, and oxidative stress (Paul, Nagano, and Robaire 2013). Highly dynamic and coordinated epigenetic processes in SSC are also vital for the fertility and health of future generations (Kosan et al. 2018). For example, several histone deacetylases (HDACs) are downregulated as adult SSCs differentiate or age (Kofman, Huszar, and Payne 2013). Globally decreased activity of the PRC2 complex was observed in the aging of SSCs (Kanatsu-Shinohara et al. 2019). Several recent genomic profiling studies have attempted to determine the landscape of epigenetic regulation in SSC (Hammoud et al. 2014; Hammoud et al. 2015; Hammoud et al. 2009; Tseng et al. 2015; Sachs et al. 2013; Kubo et al. 2015; Gan et al. 2013). However, the understanding of how aberrant gene expression patterns, epigenetic alteration, and their interaction underlie SSC aging is limited.

The maintenance of SSCs is tightly coordinated through transcriptional and epigenetic programs. In this study, we investigated how aging contributes to the deterioration of SSC function from the perspective of transcriptional and epigenetic regulations. We first established the transcriptional dynamics during SSC differentiation and delineated the stem cell intrinsic molecular program underlying the age-dependent changes in SSC function. We further established global histone modification maps of SSCs and identified the relationship between the epigenetic programming and gene expression signatures associated with SSC differentiation and aging. In addition, we mapped the dynamics of 5mC and 5hmC at single-base resolution in aged SSC and uncovered aging-associated alterations in DNA modification. At last, we examined the impact of aging on chromatin accessibility in SSC. Taken together, our study will aid in better understanding of fundamental processes in SSC aging.

## RESULTS

### Isolation and transcriptome profiling of undifferentiated and differentiating SSCs from adult testis

The common cell surface markers used for SSC isolation are also expressed in other testicular cell types to some extent and genes expressed at high levels in contaminating cells can impact RNA-seq profile interpretations (Kanatsu-Shinohara, Toyokuni, and Shinohara 2004). To overcome this problem, we utilized Oct4-GFP transgenic mice to avoid the contamination of other cell types. We first confirmed Oct4-GFP+/KIT-cells are As (single) spermatogonia or Apr (paired) spermatogonia in 3-month-old adult testis through whole-mount immunostaining (Figure 1A). In addition, FACS-sorted Oct4-GFP+/KIT-cells from adult testis possessed the capacity for *in vitro* proliferation and colony formation (Figure 1B). Based on these findings, we separated stem cells (Oct4-GFP+/KIT-) and their immediate proliferating daughter cells (Oct4-GFP+/KIT+) using FACS from neonatal (PND5.5), adult (6-8 months) and aged mice (15-18 months), which were subjected to RNA-Seq analysis (Figure 1C). Gene expression profiles of KIT- and KIT+ cells are well separated from each other in adult spermatogonia regardless of age, reflecting that they acquired more drastic transcription dynamics upon differentiation compared to the neonatal stage (Figure 1D). In contrast, neonatal KIT- and KIT+ cells clustered closely together, indicating that early germ cells share similar gene expression properties despite different differentiation status (Figure 1D). In addition, we found considerable regulatory differences between neonatal and adult spermatogonia.

**Figure 1.**
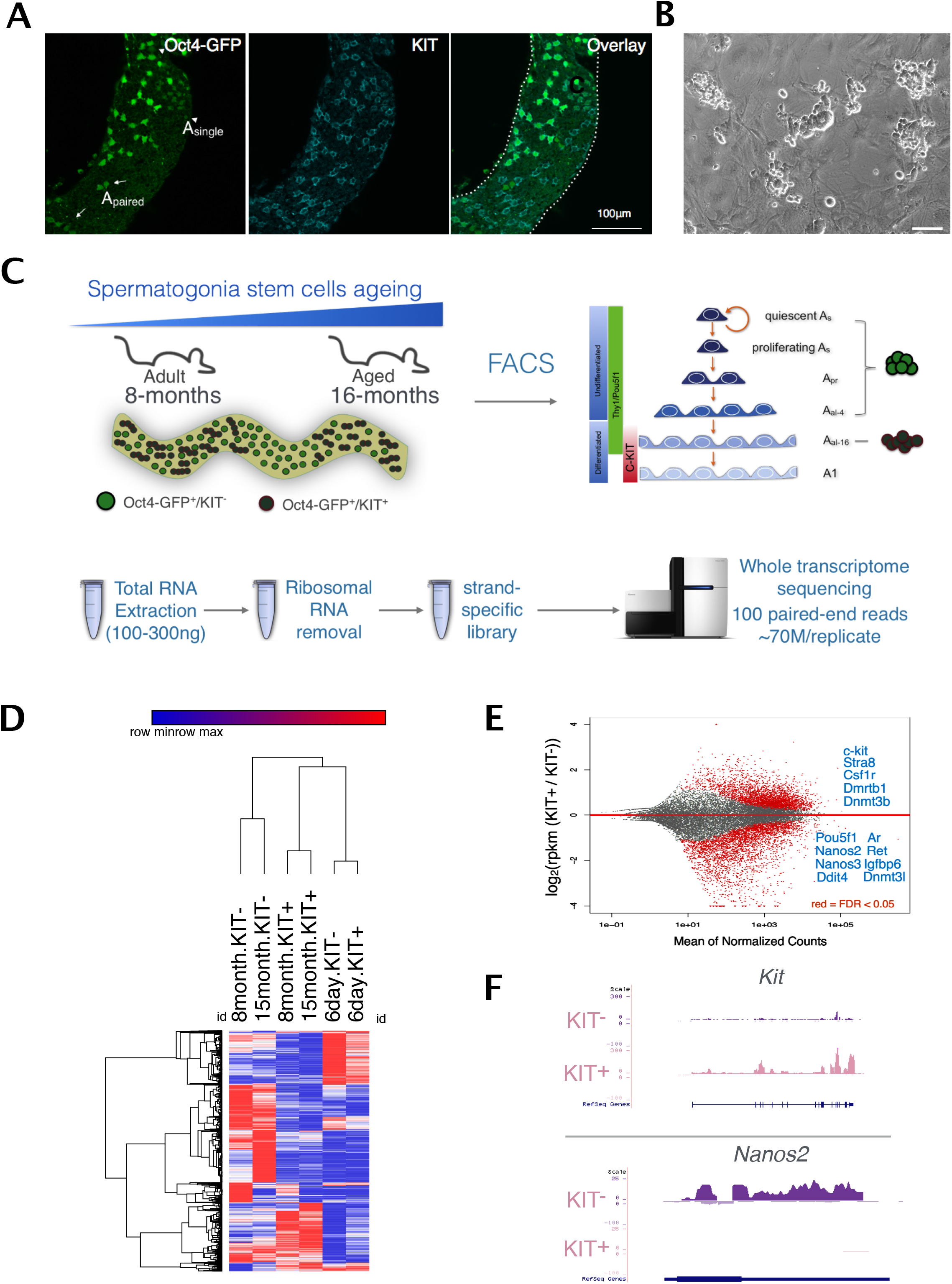
Gene expression dynamics in SSCs during the KIT transition and aging. **(A)** Whole-mount immunostaining showing the distribution of Oct4-GFP+ cells in adult testis. **(B)** Oct4-GFP+/KIT-cells isolated from adult testis can be maintained in vitro. **(C)** The schematic diagram showing the design of the sample collection and RNA-Seq analysis. **(D)** Heatmap showing global gene expression profiles of SSC populations differ significantly at different ages. **(E)** Differential gene expression between KIT- and KIT+ SSC populations in adult mice. Red dots represent differentially expressed genes with the statistical significance of FDR<0.05. Selected genes important for spermatogenesis in each fraction are shown in blue. **(F)** Genome browser view of the most significant differential gene expression, Nanos2 and Kit in KIT- and KIT+ population respectively.

### Identification of genes and pathways associated with the maintenance and aging of adult SSCs

We then compared KIT- and KIT+ cells with special focus on adult stage. The genes with higher expression level in adult KIT-comprised the genes essential for SSC selfrenewal and maintenance such as *Oct4, Ddit4, Ret* and *Nanos2*. Genes facilitating SSC differentiation were upregulated in adult KIT+ cells, such as *Stra8* and *Dnmt3b* (Figure 1E and F). IPA analysis revealed downregulation of the Mouse Embryonic Stem Cell Pluripotency pathway during differentiation (Figure S1). These results verified known regulators of SSC. Besides known essential genes for SSC self-renewal such as *Oct4* and *Id4* in this pathway, the Wnt signaling component Frizzled receptor 2 (*Fzd2*) is also upregulated in KIT-cells. Fzd2 has been reported as a marker for crypt base columnar (CBC) stem cells (Flanagan et al. 2015) and may have a role in SSC maintenance (Figure S1).

We then investigated the altered gene expression upon SSC aging in undifferentiated KIT-SSCs. IPA analysis revealed mitochondrial dysfunction and peroxisome proliferator-activated receptors (PPARs) activation as the “top targets of toxicity” as the effect of aging (Figure 2A). PPARs are important regulators in various age-associated pathophysiological processes related to energy metabolism and oxidative stress (Erol 2007), and their deregulated activation might be the culprits for the SSC aging process.

**FIgure 2.**
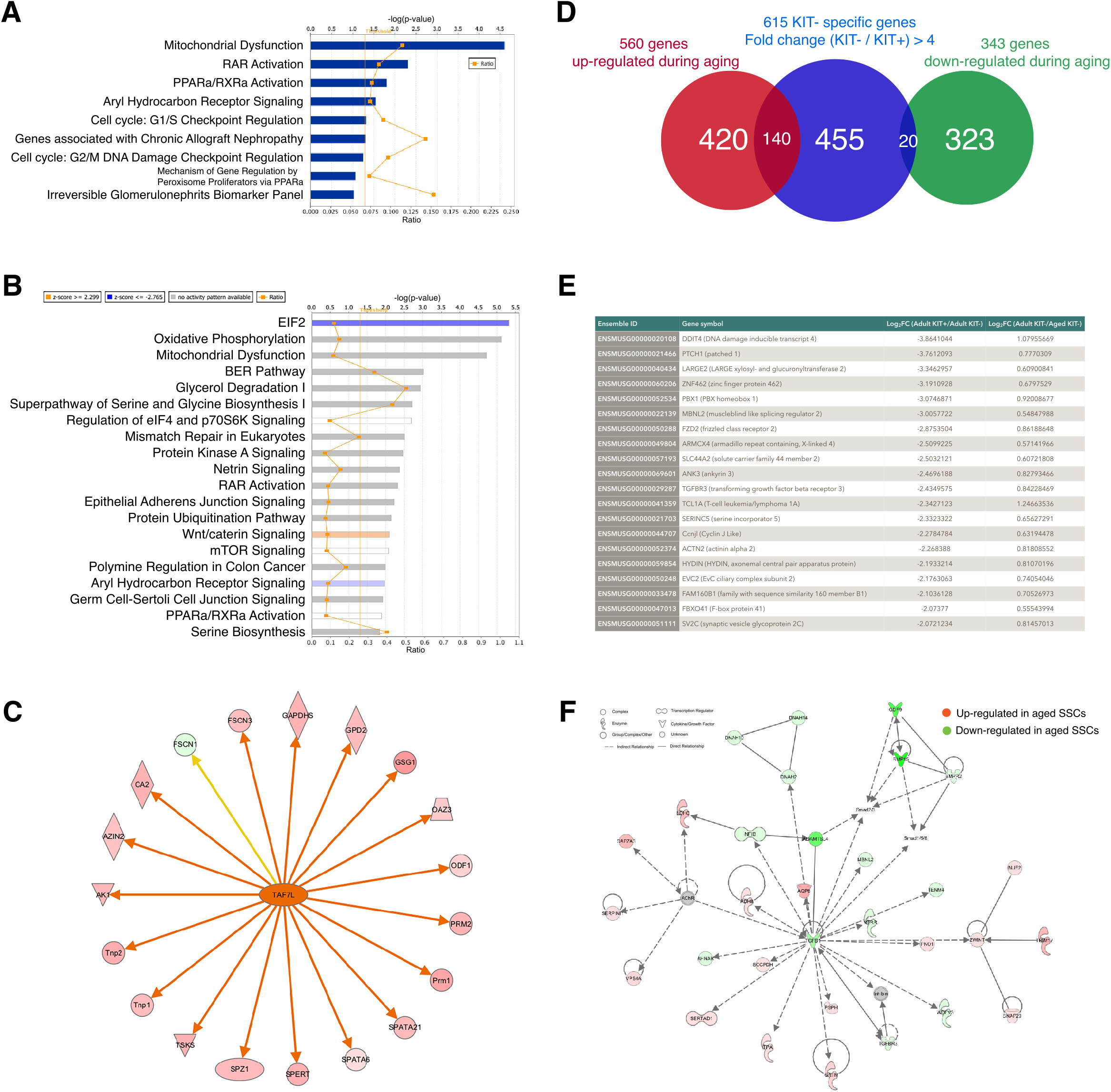
Genes and pathways associated with the maintenance and aging of adult SSCs. **(A)** Ingenuity pathway analysis (IPA) gene ontology analysis on the differentially expressed genes between adult and aged SSC showing Top toxicity target lists identified. **(B)** Top 20 canonical pathways as revealed by Ingenuity pathway analysis (IPA) gene ontology analysis on the differentially expressed genes between adult and aged SSC. **(C)** Inference of Taf7l regulation from the aging SSC transcriptome. Upstream regulator analysis by Ingenuity pathway analysis (IPA). Data illustrate Taf7l as “activated” in aged cells in upstream regulator analysis. Genes in red were greater expressed in aged cells compared with young adults, whereas genes in green were lower expressed. An orange line indicates predicted upregulation, whereas the yellow line indicates expression being contradictory to the prediction. **(D)** Aging-associated differential expression of the KIT-specific genes. Venn diagram of the overlap between genes differentially expressed upon aging and genes highly enriched in KIT-cells. **(E)** The full list of the KIT-specific genes showing down-regulation upon aging. **(F)** Inference of TGF-β signaling from the aging SSC transcriptome. Shown is the network from Ingenuity Pathway Analysis (IPA) consisting of TGF-β1 regulators (e.g., TGFB1, TGFBR3, and BMP5) and the subset of SSC aging differentially expressed genes. Symbolic representations of genes, expression changes, and regulatory relationships are shown on the top.

This is consistent with the recent finding showing decreased mitochondria numbers and expression of *Ppargc1a*, a co-activator of PPARs, in cell-intrinsic mode of SSC aging (Kanatsu-Shinohara et al. 2019). IPA canonical pathway analysis revealed eIF2 pathway, a master regulator of cell adaptation to various forms of stress, was inhibited upon SSC aging (Figure 2B), which could make SSC more susceptible to stress. Ribosome pathway was also altered, which is consistent with the previous finding that the rate of protein synthesis declines with age (Rattan 1996). Therefore, our results suggested that many common signaling pathways contributing to aging can be recapitulated in aged SSCs.

We further uncovered changes related to several spermatogonia-specific pathways that might contribute to aging. Genes with decreased expression are involved in receptor binding, cell adhesion molecule binding, integrin binding, adhesion junction and focal adhesion, suggesting that the adhesion strength between SSCs and their niche is decreased upon aging and eventually affects their long-term self-renewal potential. Strikingly, we found the aberrant expression of a large number of genes associated with spermatogenesis. For example, increased expression of the known germ cell differentiation and meiosis markers such as *Sycp2*, *Sycp3* and *Stra8* were observed in aged SSCs. Consistent with this, we found that genes upregulated in the aged cells were significantly enriched for meiosis pathway and post-meiotic development such as sperm flagellum and spermatid differentiation. IPA analysis identified *Taf7l*, which regulates a group of genes related to meiosis, as the most significant upstream regulator responsible for transcriptional changes in aging SSCs, and is required for meiotic cell cycle progression in mouse spermatogenesis (Figure 2C) (Cheng et al. 2007; Zhou et al. 2013). Interestingly, a point mutation of human *TAF7L* is associated with infertility (Akinloye et al. 2007).

We then focused on the alteration of genes preferentially expressed in KIT-cells to ask how aging affects genes presumably important for SSC maintenance. Surprisingly, we only found 20 out of 615 (3%) DEGs showed a declined trend during aging (Figure 2D). Among these genes, *Ddit4* is of particular interest because of its high expression, large fold-changes and strong specificity in KIT-cells (Figure 2E). *Ddit4* has been identified as a putative PLZF target and plays a critical role in the regulation of SSC self-renewal via inhibiting mTORC1 activity (Hobbs et al. 2010). Other notable downregulated genes include: *Mbnl2*, a member of the muscleblind protein family that modulates alternative splicing of pre-mRNAs (Wang et al. 2012); *Ptch1*, a component of hedgehog signaling which is linked with age-related diseases (Dashti, Peppelenbosch, and Rezaee 2012); *Tcl1*, a known downstream target of *Oct4*. On the other hand, 140 of KIT-enriched genes (22%) increased upon SSC aging (Figure 2D). GO analysis concurs with aforementioned results that they are mainly associated with the spermatogenesis-related function such as spermatid development, flagellated sperm motility, and fertilization.

Intriguingly, we performed network analysis using IPA to define the functional networks of aging associated genes and identify a top ranked network centred around the TGF-β pathway (Figure 2F), supporting the notion that irregularity of TGF-β signaling may contribute to the functional age-related declines that occur in stem cells with age (Sun et al. 2014).

### Alternative splicing regulation in the maintenance and aging of adult SSCs

Recent results highlight that alternative splicing also contributed to cell- and species-specific differentiation (Barbosa-Morais et al. 2012; Kalsotra and Cooper 2011). Our analyses showed that splicing regulator *Mbnl2* is highly enriched in KIT-cells and displays aberrant expression upon aging (Figure 2E). GO analysis using both age-upregulated and age-downregulated genes revealed that the major pathways associated with older SSCs were mainly involved in the processing of the primary RNA transcripts (GO:0003723: RNA binding, p<7.026E-5 and p<3.731E-17, respectively). To comprehensively analyze the alteration in the splicing landscape in SSC, we monitored the expression of all known RNA splicing regulators expressed in spermatogonia (n=143) during the KIT transition and aging. We found 51 and 20 regulators exhibited altered expression in KIT transition and during aging respectively (Figure 3A-C). For example, we observed downregulation of both *Mbnl1* and *Mbnl2* encoding muscleblind proteins and the upregulation of *Elavl2*, *Elavl3*, and *Mbnl3* during KIT transition.

**Figure 3.**
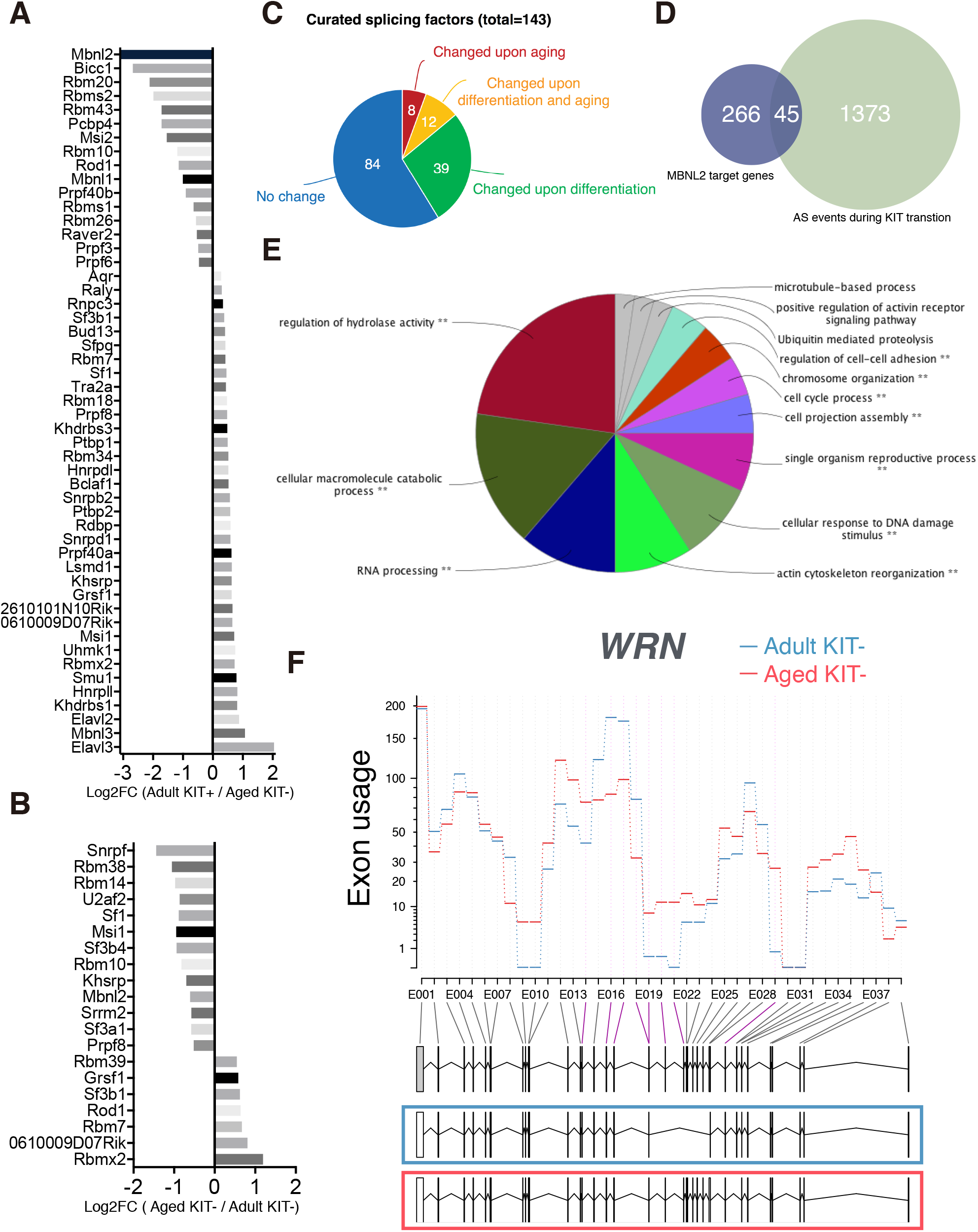
Post-transcriptional regulation during SSC differentiation and aging. **(A-B)** Column bar chart shows the fold change of splicing factors identified as differentially expressed during differentiation **(A)** and aging **(B)**. **(C)** Pie chart showing the proportion of splicing factors whose expression levels were altered during SSC differentiation and aging. **(D)** Venn diagram of the overlap between MBNL2 target genes and differential alternative splicing events detected during differentiation. **(E)** GO enrichment analysis of the altered genes by comparing adult and aged undifferentiated SSCs. **(F)** The example of WRN showed differential exon usage upon aging, indicating regulated isoform expression level at different stages.

To monitor the splicing regulation, we analyzed the differential usage of exons between KIT- and KIT+ cells using DEXseq and found 1418 genes showed evidence of differential exon usage. The alternative splicing events detected cover 45 out of 313 MBNL2 binding targets, which is consistent with the fact that MBNL2 is the most specific splicing factor enriched in KIT-cells (Figure 3D) (Han et al. 2013). On the other hand, genes with alternative splicing changes occurring during aging are associated with regulation of hydrolase activity, RNA processing, cellular response to DNA damage and cell cycle regulation (Figure 3E). We used ToppGene to survey the mouse phenotype database and found that mice with alteration in these genes shown phenotypes related to infertility such as small testis (p=1.818E-5), decreased male germ cell number (p=2.910E-5) and abnormal male germ cell morphology (p=8.766E-5). Interestingly, the Werner syndrome protein gene (*WRN*), a member of RecQ family of DNA helicases, showed differential alternative splicing (Figure 3F). Mutations in the *WRN* gene give rise to Werner syndrome (WS) and the affected individuals exhibit features of accelerated aging (Kudlow, Kennedy, and Monnat 2007).

These findings suggest transcript isoform regulation is an unexpectedly abundant regulatory mechanism in SSC differentiation and aging.

### Dynamic lncRNA expression during SSCs differentiation and aging

Recently, lncRNAs have been linked to stem cell self-renew and detrimental pathways regulating the aging process in skeletal muscle stem cells, hematopoietic stem cells and gut epithelium (Sousa-Franco et al. 2019). Therefore, we tested the hypothesis that the manifestations of aging in SSCs are associated with dysregulation of lncRNA expressions. We first employed the Ensembl annotation to provide a standardized and up-to-date analysis of lncRNA gene expression according to the biotype annotations. Out of the 3,506 genes annotated as “lincRNA”, “anti-sense” or “non-coding” (referred to as “lncRNA” for the remainder of this study), we detected 1,433 expressed at ≥ 1 FPKM.

We subsequently established a bioinformatic pipeline for detecting novel lncRNAs (See Method, Figure S2A, B). Figure S3 showed several examples of the most abundant novel lncRNAs identified. Using DESeq2, we conducted the differential gene expression analysis and performed clustering using all differentially expressed lncRNAs. Our analysis showed that lncRNAs, similar to their coding counterparts, distinguished adult from neonatal, KIT-cells from KIT+ cells and young adult from aged adult samples (Figure 4A), suggesting that the expression of lncRNAs is also tightly regulated during SSC differentiation and aging.

**Figure 4.**
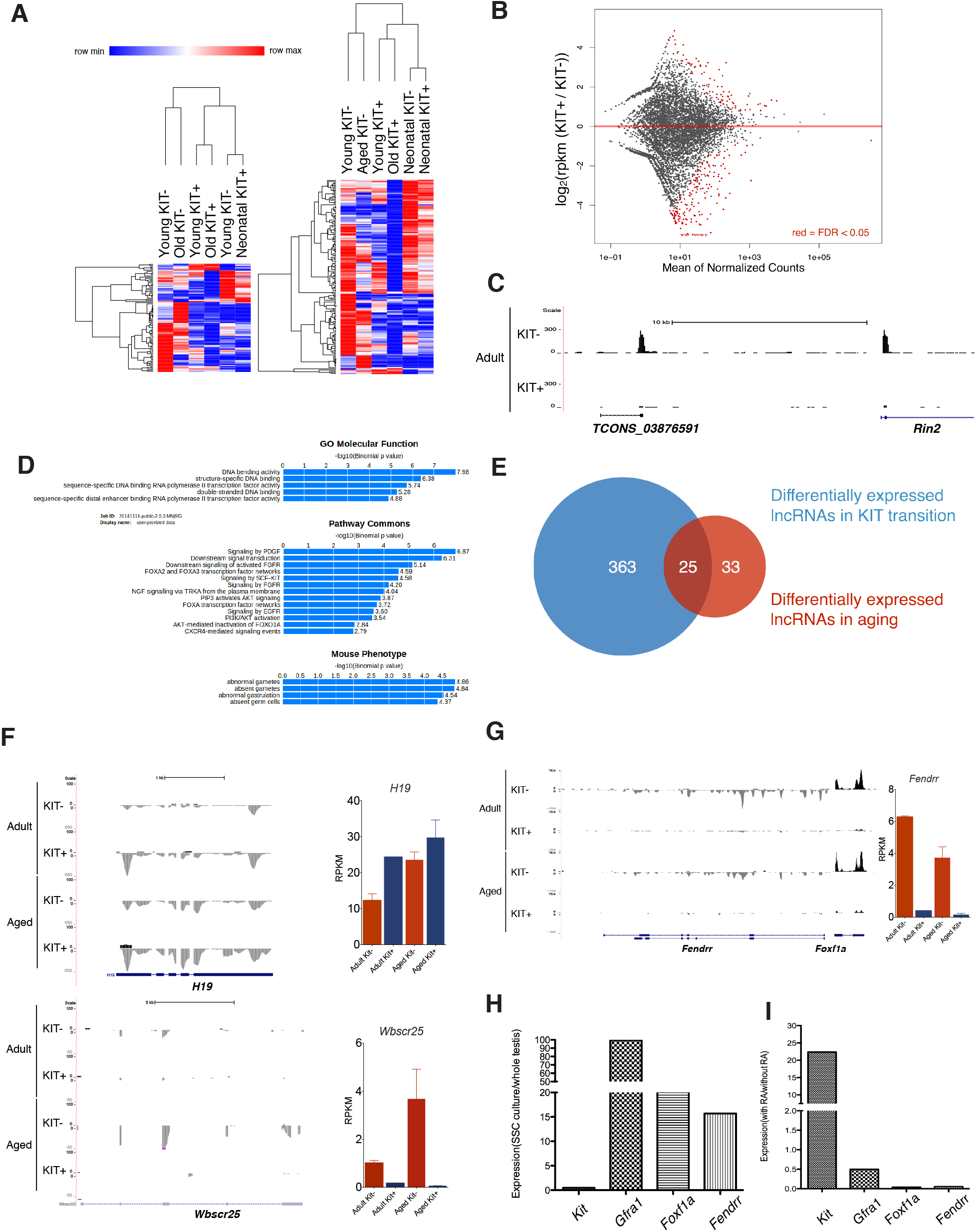
Expression of lncRNAs is tightly regulated during SSC differentiation and aging. **(A)** Heatmaps showing hierarchical clustering of differentially expressed transcripts of both known and novel lincRNA. **(B)** The MA plot shows the logarithm of the ratio between expression levels in KIT- and KIT+ populations versus the average expression of individual genes. The red dots depict the differentially expressed lncRNA (FDR< 0.05). **(C)** UCSC Genome Browser tracks for the KIT-cells specific novel lncRNA TCONS_03876591. **(D)** Functional prediction of lncRNAs regulated during differentiation by “Guilt-by-association” (GBA) analysis. **(E)** Venn diagram showing the number of lncRNAs with highly significant expression difference during KIT transition and aging (FDR<0.05) and 25 lncRNAs found commonly involved in both processes. **(F)** UCSC Genome Browser tracks for H19 and Wbscr25 with expression levels shown on the right. **(G)** UCSC Genome Browser tracks for Fendrr with expression levels shown on the right. **(H)** Relative expression levels of Fendrr and Foxf1a between SSC culture and whole testis. **(I)** Expression level changes of Fendrr and Foxf1a after RA treatment.

Next, we characterized the expression of lncRNAs within the spermatogonial differentiation paradigm. We compared the adult KIT- and KIT+ samples and identified 388 lncRNAs displaying a significant change (FDR < 0.05) (Figure 4B). For example, TCONS_03876591 is a novel lincRNA that is specifically expressed in KIT-cells (Figure 4C). It is located near *Rin2*, which was also significantly differentially expressed between KIT- and KIT+ cells. Considering that lncRNAs have been shown to regulate proximal coding genes in *cis*, we searched for protein-coding genes 1000 kb upstream and downstream of the lncRNAs (Marchese, Raimondi, and Huarte 2017). GO analysis of *cis* lncRNA targets demonstrated significant overrepresentation in terms involved in the regulation of gene expression, such as DNA bending activity, structure-specific DNA binding, sequence-specific DNA binding RNA polymerase II transcription factor activity and double-stranded DNA binding (Figure 4D), which is in line with the reported roles of lncRNA as transcriptional regulator. Pathway analysis showed that these *cis* target genes of lncRNAs were enriched in pathways that were related to spermatogenesis such as PDGF, FGF and SCF-KIT signaling pathways (Figure 4D). These findings suggested that lncRNAs might act on its neighbouring protein-coding genes in *cis* to regulate spermatogonial differentiation.

Next, we identified 58 lncRNAs with altered expression in the course of SSC aging, 25 of which are also differentially expressed during SSC differentiation (Figure 4E). For example, the lncRNA *H19* has been shown to play a role in growth, proliferation, cell cycle, apoptosis, and aging (Grammatikakis et al. 2014). Enhanced expression of *H19* due to loss of imprinting of the *H19* locus was observed in normal human prostate tissues during aging. Consistent with this, we found expression level of *H19* was elevated in aged SSCs (Figure 4F). Another candidate is *Wbscr25* (Williams Beuren syndrome chromosome region 25), which is highly enriched in KIT-cells and increased expression in aged cells (Figure 4F).

Among the known lncRNAs with significant expression changes in the process of KIT transition and aging, *Fendrr* (Fetal-lethal noncoding developmental regulatory RNA) caught our attention (Figure 4G). It displayed exclusive expression in KIT-cells, suggesting its potential role in stem cell maintenance. *Fendrr* is divergently transcribed from the transcription factor-coding gene *Foxf1a*, and they share similar expression pattern. We speculated that it should be highly expressed in SSC cultures, but at low levels in the adult testis, where true stem cells are a rare subpopulation. Using RT-PCR, we confirmed the relative expression levels of *Fendrr* and its neighbouring gene *Foxf1a* were higher in SSC cultures than whole adult testis (Figure 4H). This lncRNA-protein gene pair also underwent significant changes in response to spermatogonial differentiation, with a decrease of almost 90% after 48 hours of RA treatment (Figure 4I).

### Decreased SSC differentiation is associated with decreased gene bivalency

To investigate the epigenetic regulatory programs behind the transcriptional changes associated with SSC differentiation and aging, we first examined how histone modifications may regulate SSC KIT transition. We performed genome-wide analysis of H3K4me3 and H3K27me3 by ChIP-Seq and identified a set of bivalent promoters (Figure 5A). We found bivalent promoters were overrepresented in genes encoding cell–cell signaling molecules, developmental regulators, cell adhesion molecules and embryonic morphogenic proteins. Previous reports have shown that bivalent promoters are dynamically regulated during stem cell differentiation, but the role of bivalency in the course of SSC differentiation remains largely unknown (Mikkelsen et al. 2007; Cui et al. 2009). Globally, we found a moderate inverse correlation between the change in the enrichment of H3K27me3 with the change in expression of differentially regulated genes during SSC differentiation (Figure 5B). A closer examination revealed a subset of bivalent promoters showed a significant decrease of the repressive H3K27me3 mark but maintained the active H3K4me3 mark, which led to an induction of gene expression (e.g. *Kit* and *Stra8*) during differentiation (Figure 5C and S4A, B). This was further validated by demonstrating the significant gene expression changes (log2FC) that were associated with the above epigenetic changes during the KIT transition. Unlike stable bivalent promoters which have a predisposition for downregulation during differentiation, genes with decreased level of H3K27me3 were associated with a significant upregulation of gene expression during the KIT transition (Figure 5D). GO analysis showed that this set of bivalent genes are associated with plasma membrane and cell junction (Figure 5E). Our data thus show that while the genomic location of H3K27me3 is mostly conserved, the extent of gene bivalency differs quantitatively during KIT transition. This demonstrates the involvement of resolving bivalency in transcriptional activation in SSC differentiation, which is in line with ESC differentiation (Voigt, Tee, and Reinberg 2013).

**Figure 5.**
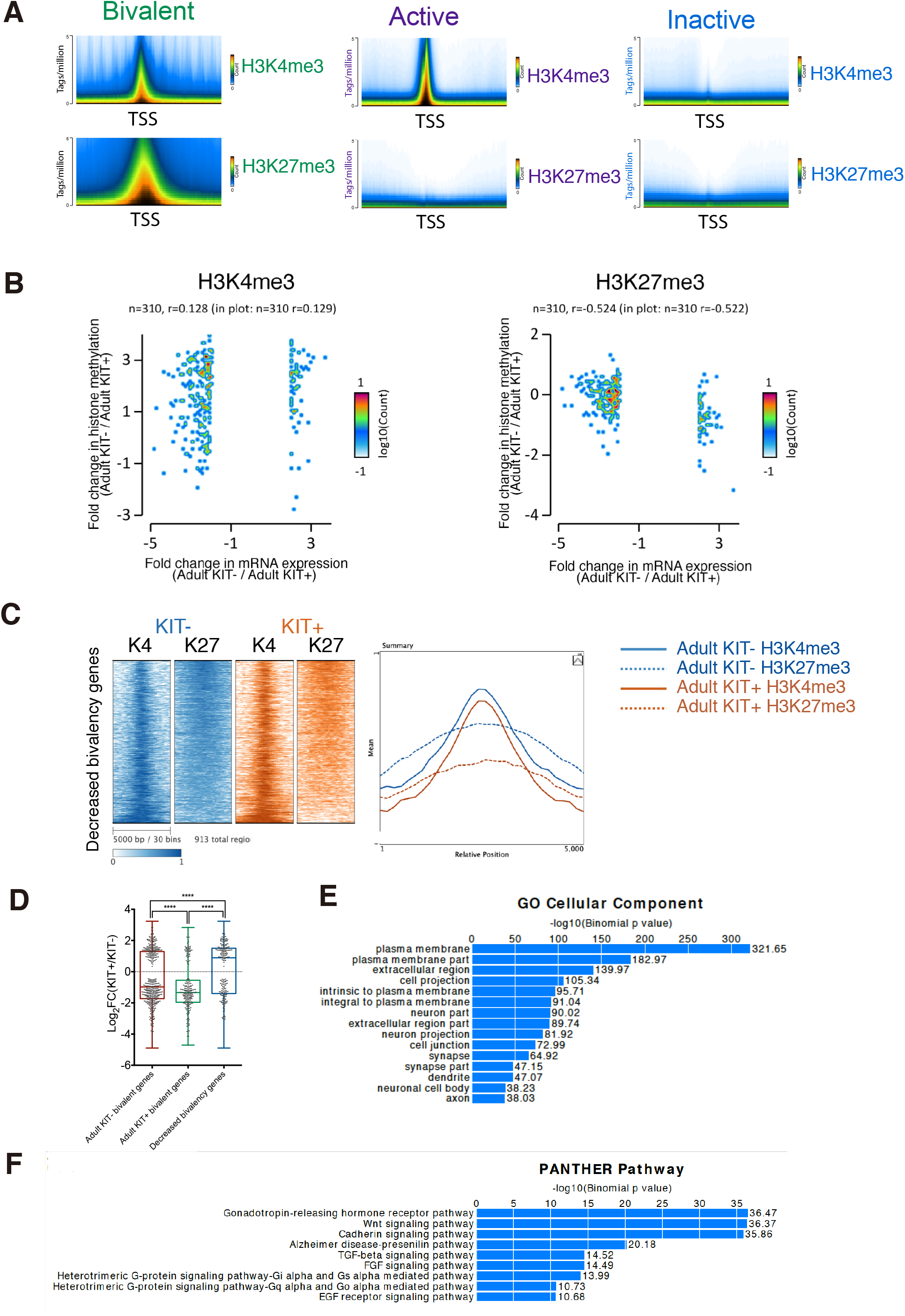
Gene bivalency on transcriptional regulation during the adult KIT transition. **(A)** The average profile plots for each category of promoter. **(B)** Negative correlation of H3K27me3 difference and gene expression fold change. **(C)** Heat maps (Left) and meta-plot (Right) of histone modifications across the 5 kb region centred at the TSSs of the genes with decreased bivalency during the KIT transition. **(D)** Box plots represent the spread of the gene expression changes as a result of the loss of bivalency during the adult KIT transition. The whiskers represent 5-95 percentile with the medians shown as horizontal lines. The p-values were calculated using unpaired, two-tailed t-test with 95% confidence. **(E)** Bar plot showing enrichment of biological processes for genes with decreased bivalency. **(F)** Bar plot showing enrichment of biological processes for genes associated with decreased H3K27me3 marks during SSC aging.

It has been suggested that PcG-mediated H3K27me3 alteration drives many age-related changes and is often dysregulated in human malignancies (Greer et al. 2010; Jones and Rando 2011; Molofsky et al. 2006). Therefore, we continued to examine the bivalent domain in aged SSCs. We directly compared the epigenetic states of RefSeq promoters in adult KIT- and aged KIT-cells and found the distributions of H3K4me3 and H3K27me3 were highly similar (Figure S4C). As a result, no significant correlation between changes in levels of mRNA and H3K4/K27me3 levels during aging were found (Figure S4D). We employed SICER to identify differentially histone methylation enrichment across the genome (Zang et al. 2009). 7080 differentially enriched H3K27me3 peaks were identified between adult KIT- and aged KIT-cells, in which 1734 peaks have increased enrichment in aged cells and 5346 peaks with reduced enrichment (Figure S5). GO analysis performed on the list of genes associated with decreased H3K27me3 mark in SSCs during aging showed the enrichment for GnRH, Wnt/β-catenin, and TGF-β pathway (Figure 5F).

### Age-dependent 5mC and 5hmC alteration in SSCs

We continued to investigate the effect of aging on DNA modification in SSCs. We performed reduced-representation bisulfite sequencing (RRBS) and compared the DNA methylation level (5mC+5hmC) in young and old SSCs. We found that the level of (5mC+5hmC) at individual CpG sites correlates well between the young and old SSCs (Figure 6A). The 5mC differences at gene promoters are extremely rare and we only identified 3 genes, *Sfi1, Slc22a2*(*Oct2*) and *Gtf2f1* with a decrease in methylation larger than 30% (covered at least by 5 CpG sites) (Figure 6B). Interestingly, both *Sfi1* and *Slc22a2* have been implicated in the aging process, as the aging-associated methylation change have been observed in several reports (Cheng et al. 2008; Park et al. 2014). Many studies showed that 5hmC is linked with aging, but the role of 5hmC in SSC aging is largely unknown (Lopez, Fernandez, and Fraga 2017). We first confirmed the active demethylation process in spermatogonia by immunostaining. Indeed, 5hmC and its oxidative products 5caC and 5fC are specifically located in the spermatogonia at the basal membrane (Figure 6C). Since conventional RRBS cannot distinguish 5mC and 5hmC, we employed oxRRBS which allowed us to measure 5mC and 5hmC separately (Booth et al. 2013). In general, the levels of total 5hmC observed across the genome are approximately 10 folds lower than those of 5mC (Figure 6D). When we compared 5mC and 5hmC level between the two groups, we noticed global 5mC was slightly but significantly decreased, while 5hmC level was increased in aged SSCs (Figure 6E). Plotting the average 5mC level across all genes revealed a global hypomethylation at the gene body in aged SSC, akin to that observed in humans (Figure 6F) (Hannum et al. 2013; Horvath 2013). The reduced 5mC level at gene body was accompanied by an increased 5hmC level (Figure 6F). Indeed, we found a very strong inverse correlation between reduction of average 5mC and increased level of 5hmC at gene bodies (Figure 6G). These results suggested that 5hmC-dependent active demethylation is associated with 5mC alteration during aging.

**Figure 6.**
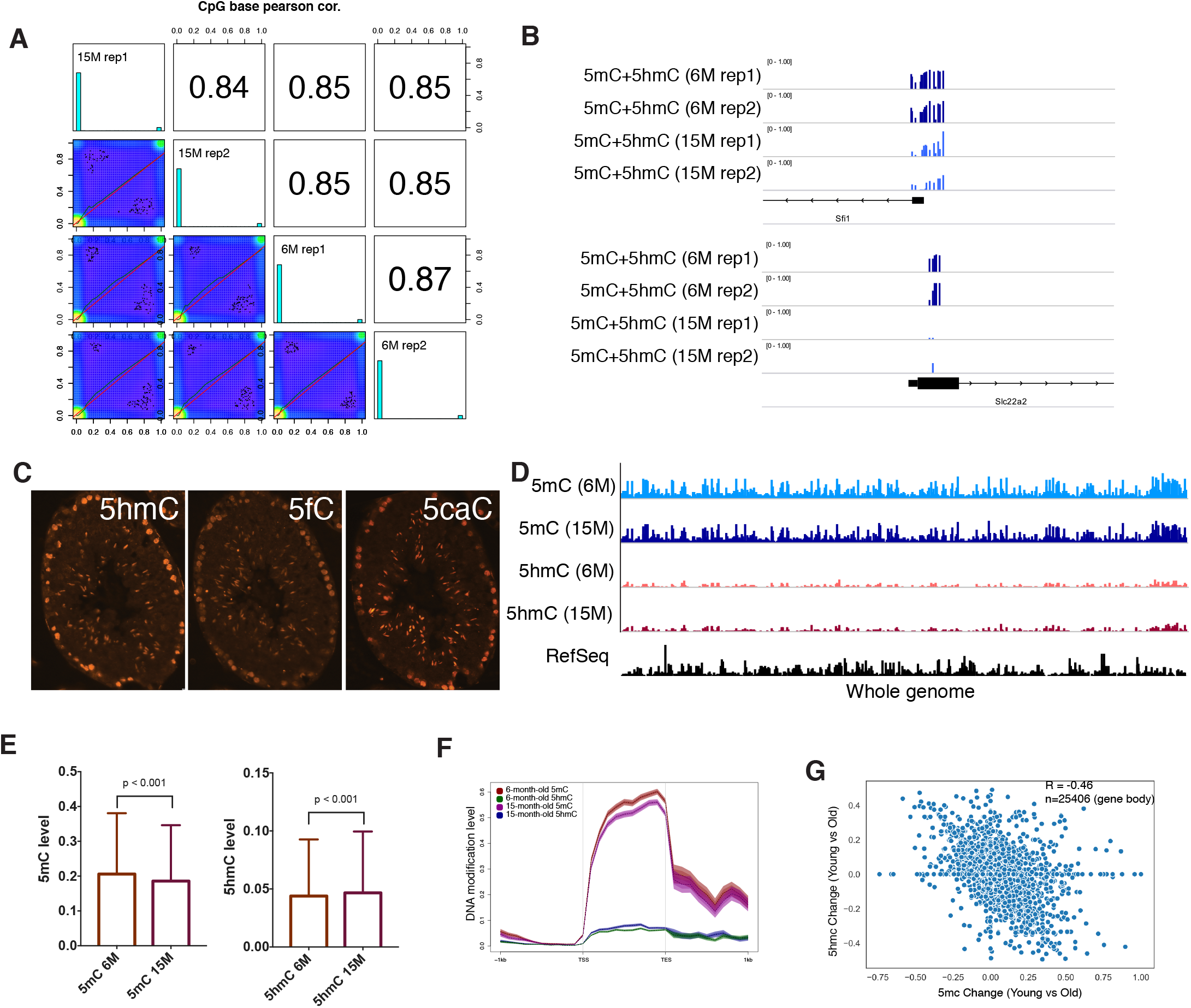
Effect of aging on 5mC and 5hmC in SSC. **(A)** Scatter plot and correlation of CpG methylation between RRBS methylomes. Scatter plots of percentage methylation values for each pair in four libraries. Numbers on the upper right corner denote pairwise Pearson’s correlation scores. The histograms on the diagonal are methylation distribution of CpG sites for each sample. **(B)** Examples of genomic RRBS profiles of differentially methylated promoters. **(C)** Representative images of testicular sections from adult mice showing immunofluorescent staining of 5hmC, 5fC, and 5caC, which were observed in cells residing on the basement membrane. **(D)** Genome browser view of 5mC and 5hmC distribution in all chromosomes. **(E)** The box plots show the average 5mC and 5hmC levels per bin (100kb) of the genome in young and aged SSCs. A pairwise comparison was performed using the independent sample t-test. **(F)** Metagene density profiles of 5mC and 5hmC for RefSeq transcripts showed levels of 5hmC are consistently higher in the aged SSCs than in the young SSCs along the length of the gene. **(G)** Scatterplot of “switching” between 5mC and 5hmC at gene bodies in aged SSC relative to young SSC.

## DISCUSSIONS

Understanding the underlying mechanisms conferring the functional decline in aged SSCs is one of the major questions to be addressed in reproductive biology. Early studies mainly focus on gene expression alteration during SSC aging (Kokkinaki et al. 2010; Paul, Nagano, and Robaire 2013). These microarray analyses lacked the specificity of sequencing-based techniques and could not reveal gene isoform and novel transcript regulation. In this regard, our study provided a global view of the transcriptional and epigenetic programming that is associated with SSC normal differentiation and aging.

Our transcriptome profiles of spermatogonia provide a comprehensive transcriptomic map and reliable resource to study the molecular mechanisms of spermatogonial differentiation and the effect of aging. Aged SSCs apparently have a significantly altered gene expression signature compared to their younger counterparts. Pathway analysis revealed that many common age-related transcriptomic changes underlying both aging and the pathogenesis of multiple age-related diseases are also reflected in aged SSCs, illustrating the robustness and relevance of the current study. We also identified spermatogonia-specific aging pathways and our most striking observation is the aberrant expression of a large number of genes associated with meiosis pathway. Network analysis suggested that *Taf7l* might be one plausible candidate underlying this regulation. One possible explanation is that the mitotic and meiotic cell cycles should be kept in a tight control in SSC, but when germ cells are driven aberrantly from the mitotic into the meiotic cell cycle, their stem cell properties are lost.

A better understanding will not be possible without unravelling the essential mechanisms involved in the maintenance of SSCs and their differentiation in steady spermatogenesis. We have identified genes showing strong enrichment in KIT-cells, which might represent those important for SSC maintenance. We then examined whether there is an age-associated loss of their expression in mouse testes. Interestingly, we only identified a small percentage of genes enriched in undifferentiated SSC with decreased expression during aging. Nevertheless, this analysis leads us to propose several SSC-specific factors including *Ddit4*, which action mainly occurs through inhibition of the mTOR pathway (Brugarolas et al. 2004). mTOR can regulate stem cell function. Functional interaction between mTOR and PLZF is a critical rheostat for maintenance of SSC self-renewal, and this action is through *Ddit4* (Hobbs et al. 2010). Hyperactive mTOR signaling has also been shown to have a plausible role involved in regulating aging in mammals (Johnson, Rabinovitch, and Kaeberlein 2013). Therefore, *Ddit4* might contribute to aging through modulating the mTOR pathway which regulates lifespan in multiple species. It has been demonstrated that *Ddit4* expression increased significantly in response to calorie restriction in both rats and mice, which is the only intervention known to extend lifespan (Linford et al. 2007; Selman et al. 2006). Altogether, this finding points to *Ddit4* as a direct link between aging pathology and stem cell function decline in SSCs.

Epigenetic integrity is an essential element for maintaining normal stem cell function during aging (Jones and Rando 2011). Aged stem cells in mouse models feature aberrant expression of chromatin-modifying enzymes in various stem cell compartments (Kamminga et al. 2006; Sun et al. 2014). Consistent with this, we also found that the expression of some chromatin modulators were ablated during SSC aging. In addition, we present here a detailed examination of H3K27me3 modification in SSC differentiation and aging. First, we found a specific set of genes encoding developmental regulators were bivalently marked with H3K4me3 and H3K27me3, which are significantly reduced in the course of differentiation. This suggested that bivalent domains maintain developmental genes in a silent state in undifferentiated cells while keeping them poised for subsequent induction upon SSC differentiation. Second, locus-specific alteration of H3K27me3 level was observed during SSC aging. It has been reported that, however, the change in the level of H3K27me3 could not be linked to significant changes in mRNA levels of the associated genes. One possible explanation is that steady-state mRNA levels detectable by RNA-seq are not an accurate reflection of active transcription. Another possibility is that SSC aging and age-related differential gene expression are multifactorial in nature and regulated by several other mechanisms. Previous characterization of SSC in vitro aging showed that H3K27me3 peaks were significantly decreased in the promoter regions of *Wnt7b* which resulted in *Wnt7b* up-regulation*(Kanatsu-Shinohara et al. 2019)*. Similarly, we found genes associated with reduced H3K27me3 were related to common aging pathways such as Wnt and TGF-β pathways and their contribution to SSC aging has yet to be elucidated.

Current literature on how aging affects DNA methylation has been controversial. One possibility is that methylation levels in tissues have been measured instead of specific cell types. Our results analyzed the purified SSCs and revealed there is a decrease in average 5mC level in gene bodies. Our results also indicated an age-related increased 5hmC level and the observed changes raised the possibility that 5hmC acts as an intermediate step of active DNA demethylation and contributed to hypomethylation. We further identified several promoters displaying changes in DNA methylation occur with age and may be functionally important. For example, one of the hypomethylated genes *Sfi1* is related to centrosome amplification, and interestingly, an age-related change in centrosome amplification was also identified in aged intestinal stem cells (Park et al. 2014). Moreover, upregulation of *Sfi1* probably causes other centrosome aberrations such as centrosome misorientation, which can lead to cell cycle arrest and consequently decline in spermatogenesis during aging (Cheng et al. 2008).

In conclusion, our study indicates that the aging process in mouse SSCs is associated with transcriptome alteration, which is connected with epigenetic changes and may be mediated via histone modification, methylation and hydroxymethylation of DNA. We envision that our results could lay the groundwork for further exploration into the influence of epigenome dynamics in human reproductive aging.

## MATERIALS AND METHODS

### Animals

All animal procedures were performed according to protocols approved by the Animal Experimentation Ethics Committee (AEEC) of the Chinese University of Hong Kong and following the Animals (Control of Experiments) Ordinance (Cap. 340) licensed from the Hong Kong Government Department of Health. Oct4-GFP transgenic mice (B6; CBA-Tg(Pou5f1-EGFP)2Mnn/J, Stock no.: 004654) were obtained from The Jackson Laboratory (Jackson Laboratory, Bar Harbor, ME, USA. Male transgenic mice at different ages (i.e., PND5.5 for neonatal group, 6-month-old for young group and 15-month-old for aged group) were employed in this study. Age and strain matched non-reporter mice i.e. C57BL/6 mice were used for cell purification using FACS sorter as negative control devoid of GFP expression.

### Cell isolation, sample preparation and sequencing

Testicular cells from adult and aged Oct4-GFP mice were isolated by a two-step enzymatic digestion. For RNA-seq experiment, total RNA was isolated from each cell fraction using the AllPrep DNA/RNA Mini kit (Qiagen, Valencia, CA, USA) and sequencing libraries were prepared with ribosomal RNA (rRNA) depletion using a Ribo-Zero Gold kit (Epicentre) followed by Apollo 324 NGS Library Prep System (WaferGen Biosystems, Fremont, CA, USA). Two biological replicates of the prepared libraries were sequenced on an Illumina HiSeq 2000 (Illumina, San Diego, CA) with 100 base pairs (bp) paired-end RNA-Seq reads. gDNA extracted from the same batch of cells are subjected to RRBS and oxRRBS experiments. 2×10^5^ freshly sorted Oct4-GFP+/KIT- or Oct4-GFP+/KIT+ cells were cross-linked with 1% formaldehyde for 10 min. ChIP experiments and ChIP-Seq libraries were prepared using Diagenode True MicroChIP kit according to the manufacturer’s protocol.

## AUTHOR CONTRIBUTIONS

JL and TL designed the study, performed experiments, analyzed and interpreted data, and prepared the manuscript. JL, HCS, ACSL and TL reviewed and edited the manuscript. All authors read and approved the final manuscript.

## ACKNOWLEDGEMENT

The authors acknowledge the support of Core laboratory in School of Biomedical Sciences, The Chinese University of Hong Kong and Prof. Bo Feng (School of Biomedical Sciences, Faculty of Medicine, The Chinese University of Hong Kong) for providing Oct4-GFP transgenic mice. This work was supported by General Research Funds from Hong Kong Research Grant Council (Project no. 469713) and Lo Kwee-Seong Biomedical Research Seed Fund from School of Biomedical Sciences, The Chinese University of Hong Kong (Project no. 7104687).

