## Supplemental methods and figure legends for "Transcriptomic and epigenomic profiling of young and aged spermatogonial stem cells reveals molecular targets regulating differentiation"

### SUPPLEMENTAL EXPERIMENTAL PROCEDURES

#### Immunohistochemistry for whole mount tubules and imaging

Seminiferous tubules were decapsulated from mouse testes and gently pulled apart with a pair of fine forceps in petri dishes. The resulting tubules were then fixed in 4% paraformaldehyde solution for 2 hours followed by permeabilization with 0.2% Triton-X and dehydration in a graded series of methanol washes. After rehydration in PBS, tubules were processed with staining protocol. Briefly, sections were permeabilized in PBS with 0.2% Triton-X and blocked with PBS supplemented with 0.2% Triton X-100 and 10% normal donkey serum (Jackson ImmunoResearch) for 1 hour at room temperature, followed by incubation with primary antibody overnight at 4 °C. After washing in PBS, sections were incubated with corresponding fluorescence-conjugated secondary antibody. Finally, the slides were washed with PBS and mounted with PermaFluor aqueous mounting medium (Thermo Fisher). Primary and secondary antibodies utilized in this study are listed in Table 1. Fluorescent microscopy and white field images were acquired using a Leica SP2 confocal microscope. All images were captured using Leica Application Suite Software.

#### Isolation of undifferentiated and differentiating spermatogonia by fluorescence-activated cell sorting (FACS)

Testicular cells from adult and aged Oct4-GFP mice were isolated by a two-step enzymatic digestion. Firstly, the isolated tubules were incubated in RPMI with 1 mg/ml type 2 collagenase (Worthington) and 5μg/ml DNase I (Sigma-Aldrich) at 37°C for 20 minutes with occasional shaking. After several washes by sedimentation in RPMI, tubular fragments were digested in RPMI with 1 mg/ml type 2 collagenase (Worthington), 1 mg/ml hyaluronidase (Sigma-Aldrich) and 5μg/ml DNase I (Sigma-Aldrich) at 37°C for 30 minutes with occasional shaking. Single-cell suspensions were obtained by passing through a cell strainer (40μm). The resulting cell suspensions were directly separated by centrifugation in 27% Percoll to concentrate spermatogonia and to reduce cellular debris. Spermatogonia-enriched bottom fraction was collected and washed followed by processing to antibody staining. After incubation with an antibody against KIT at 4°C for 30 minutes, cell fractions were collected with a FACS Aria II cell sorter (BD Biosciences). Age and strain matched non-reporter mice i.e. C57BL/6 mice were used for cell purification as negative control devoid of GFP expression for cell sorting experiments. Positive antibody labeling was determined by comparison to staining with isotype control antibodies.

#### RNA-Seq

1x10^6^ primary cells of each population in two independent experiments were FACS-sorted as described above. Total RNA was isolated from each cell fraction using the AllPrep DNA/RNA Mini kit (Qiagen, Valencia, CA, USA) according to the manufacturer's instructions. RNA yield and purity were evaluated using a NanoDrop 1000 spectrophotometer (NanoDrop Technologies/Thermo Scientific, Waltham, MA). Quality and integrity were determined by RNA 6000 Nano kit with a Bioanalyzer 2100 (Agilent Technologies, Santa Clara, CA). The ratio of absorbance at 260 and 280 nm was ≥1.9 and the RNA integrity number (RIN) was >9 for all samples. Sequencing libraries were prepared with ribosomal RNA (rRNA) depletion using a Ribo-Zero Gold kit (Epicentre) followed by Apollo 324 NGS Library Prep System (WaferGen Biosystems, Fremont, CA, USA). Two biological replicates of the prepared libraries were sequenced on an Illumina HiSeq 2000 (Illumina, San Diego, CA) with 100 base pairs (bp) paired-end RNA-Seq reads.

#### Data analysis of bulk RNA-Seq experiments

Raw reads were pre-processed with Trimmomatic for Illumina adaptor sequences trimming and mapped to mouse (mm9) genome. Counts were assigned to genes defined by the Ensembl (release 67) annotation using featureCount R function. Differentially expressed genes were identified with DESeq2 package (Love et al., 2014) and the differential exon usage analysis was done using DEXSeq (Anders et al., 2012).

#### Functional Annotation of Differentially Expressed Genes by IPA

To define the functional networks of differentially expressed genes, data was analyzed by using the Ingenuity Pathway Analysis (IPA, Ingenuity Systems) that calculates a significance score (network score) for each network. A data set containing the gene identifiers and their corresponding fold change (log2) values were uploaded into the IPA software. Each gene identifier was mapped to its corresponding gene object in the Ingenuity Pathways Knowledge Base to identify molecules whose expression was significantly differentially regulated (focus genes or Networks Eligible molecules). These focus genes were overlaid onto a global molecular network developed from information contained in the Ingenuity Knowledge Base. Networks of these focus genes were then algorithmically generated based on their connectivity. A network is a graphical representation of the molecular relationships between genes or gene products, which are represented as nodes, and the biological relationship between two nodes is represented as an edge (line). All edges are supported by at least one reference from the literature, or from canonical information stored in the Ingenuity Pathways Knowledge Base.

#### Identification and quantification of lncRNA

As a basis for our analysis of lncRNA expression, we utilized our high quality, ribosome RNA-depleted RNA-Seq dataset. We also included poly(A)+ RNA-Seq dataset obtained from THY1+/KIT- cells that were isolated from adult mice (Hammoud et al., 2014). and additional two RNA-Seq datasets obtained from BSA gradient enriched neonatal spermatogonia (Gan et al., 2013; Soumillon et al., 2013). Together, we combined a total of ~1 billion uniquely mapping sequencing reads to survey lncRNA expression in SSC (Figure S3A).

We employed *ab initio* transcript assembly to assess the completeness of the known annotation about lncRNAs expressed in SSCs. Raw reads were mapped to the mouse mm9 reference genome using TopHat and *de novo* transcriptome assembly of mapped reads was performed using Cufflinks. Gencode, Noncode and RefSeq annotated transcripts were merged into one set of gene annotation, and these annotated transcripts were filtered out from Cufflinks-assembled transcriptomes. The novel transcripts were then categorized into different categories according to their locations compared with the reference genes. To exclude artifacts of the sequence alignment process, unspliced intronic pre-mRNA, and genomic DNA contamination, only lncRNAs transcribed within intergenic and non-exon-overlapping regions were retained. We further filtered the list of novel transcripts by CAGE signaling, RNA PolII binding and H3K4me3 enrichment, and determined the Coding Potential Calculator (CPC) score of each transcript with default setting. Finally, overlapping lncRNA loci were merged to create a single 'consensus' transcript model.

#### ChIP-Seq library preparation and sequencing

ChIP experiments and ChIP-Seq libraries were prepared using Diagenode True MicroChIP kit according to the manufacturer's protocol. 2x10^5^ freshly sorted Oct4-GFP+/KIT- or Oct4-GFP+/KIT+ cells were cross-linked with 1% formaldehyde for 10 minutes. The crosslinking was stopped by incubation with 0.125M glycine at room temperature for 5 minutes. Chromatin was fragmented to 200 to 600 bp by sonication using a Bioruptor sonicator (Diagenode, Liège, Belgium). 1ug antibodies against H3K4me3 or H3K27me3 were used for ChIP. Half of the chromatin was used for each IP, and 10% of the chromatin solution was reserved as input control. The chromatin immunoprecipitated (ChIPed) DNA was subjected to library preparation using MicroPlex Library Preparation kit (Diagenode, Liège, Belgium). The prepared library was sequenced with single-end 50bp reads using the Illumina HiSeq 2000.

#### ChIP-Seq data analysis

Reads were mapped to the mouse genome (mm9 assembly) using Bowtie (Langmead et al., 2009). Mapped reads were filtered for PCR duplicates using MACS2 (Zhang et al., 2008). Peak calling and visualization of tracks were done using EaSeq (Lerdrup et al., 2016) and heat maps were made using EaSeq or seqMINER (Ye et al., 2011). In our annotation, a TSS region is defined as ±2.5 kb relative to the TSS. For density plots, we generated bigwig files allowing only one read per chromosomal position, eliminating potential spurious spikes. For density plots, the regions of ±5 kb for all annotated TSSs were divided into 100 equal-sized bins.

#### Reduced representation bisulfite sequencing (RRBS) and oxRRBS

RRBS libraries from oxidized and non-oxidized DNA were prepared using TrueMethyl Kit according to the manual's instruction (Cambridge Epigenetix). Briefly, 0.5ug of genomic DNA was digested with MspI followed by end repair, A-tailing, and ligation of adaptors. Adaptor-ligated MspI-digested DNA was size-selected (110-380 bp) and purified. 1/5 the DNA was kept for the generation of the non-oxidized library and remaining DNA was subjected to oxidation. Both oxidized and non-oxidized DNA samples were bisulfite-treated. Final library amplification was carried out for 18 cycles, after which the libraries were purified using AMPure XP beads (Agencourt) and sequenced on Illumina platform. The raw reads were quality-checked with FastQC (http://www.bioinformatics.babraham.ac.uk/projects/fastqc/), and low-quality reads and adapters were removed using Trim Galore (http://www.bioinformatics.babraham.ac.uk/projects/trim_galore/) using '--rrbs' trimming mode. Trimmed sequences were mapped to the mouse genome with Bismark (Krueger and Andrews, 2011). The extracted methylation calls were subjected to the subsequent in-depth analysis performed with R packages (http://www.r-project.org/) or customized scripts.

#### Germ cell culture

Oct4-GFP+/KIT- cells were isolated from 3-month-old Oct4-GFP mice by FACS cell sorting as described above. Cells were plated at a density of 1.5 to 2 x 10^5^ per well on 12-well plates with mitomycin treated (Sigma-Aldrich) MEFs feeder layers and cultured in a serum-free medium consisting of StemPro-34 SFM medium supplemented with StemPro-34 nutrient supplement (Life Technologies, Carlsbad, CA, USA), 0.2% bovine serum albumin (MP Biochemicals, Santa Ana, CA), 1% fetal bovine serum (embryonic stem cell-qualified, Life Technologies), 50 U/ml penicillin-streptomycin (Life Technologies), 2 mM GlutaMAX (Life Technologies), 1% mM non-essential amino acids (Life Technologies), 1% minimal essential medium (MEM) vitamin solution (Life Technologies), 1 mM sodium pyruvate (Life Technologies), 50 μM 2-mercaptoethanol (Sigma-Aldrich), 25 μg/ml insulin (Sigma-Aldrich), 100 μg/ml transferrin (Sigma-Aldrich), 60 μM putrescine (Sigma-Aldrich), 30 nM sodium selenite (Sigma-Aldrich), 1 mg/ml D-(+)-glucose (Sigma-Aldrich), 1 μl/ml Dl-Lactic acid (Sigma-Aldrich), 60 ng/ml progesterone (Sigma-Aldrich), 30ng/ml β-estradiol (Sigma-Aldrich), 10 μg/ml D-biotin (Sigma-Aldrich), 100 μM ascorbic acid (Sigma-Aldrich), 40 ng/ml human GDNF (R&D Systems, Minneapolis, MN, USA), 20 ng/ml mouse epidermal growth factor (Life Technologies), and 10ng/ml human basic fibroblast growth factor (BD Biosciences, San Jose, CA, USA). Cultures were maintained at 37°C in an incubator with humidified 5% CO2 and 95% air atmosphere. Half of the medium was replaced with fresh medium every 2–3 days and cells were passaged enzymatically using 0.25% trypsin/EDTA (Life Technologies) at a ratio of 1: 2 or 1: 3 every 6 to 7 days onto new mitomycin-treated MEF plates. For RA-induced differentiation, 5mM all-trans-RA (Sigma-Aldrich) stock in ethanol was diluted to 1 μM in medium before applying to cells while vehicle (0.1% ethanol) was added in the control group.

#### Gene Ontology (GO) analysis

GO enrichment was performed using DAVID (Huang da et al., 2009), ToppGene Suite (Chen et al., 2009). A hypergeometric test with the Benjamini and Hochberg false discovery rate (FDR) was performed using the default parameters to adjust p-value.

### SUPPLEMENTAL FIGURES LEGENDS

#### Figure S1. Schematic representation of the Mouse Embryonic Stem Cell Pluripotency pathway highlighting the members identified as preferentially expressed in KIT- cells.

Genes highlighted in red are identified as preferentially expressed in KIT- cells while genes labeled in green show higher expression in KIT+ samples. Noted that As marker *Id4* is identified in this analysis.

#### Figure S2. Detection and quantification of lncRNAs

(A) Data sources for novel lncRNA identification. (B) A bioinformatics pipeline for discovery lncRNAs in SSC. See text and Methods session for details. Raw reads are first mapped onto the reference mouse genome. The initial assemblies are categorized by cuffcompare, compared with the combined gene annotations. The lncRScan program is performed to detect the novel lncRNAs from the high-quality assemblies according to multiple criteria.

#### Figure S3. UCSC Genome Browser tracks for the most expressed novel lncRNAs.

#### Figure S4. Analysis of histone modification dynamics in SSC aging.

(A) and (B) Genome browser representation of H3K4me3 and H3K27me3 modification at selected genes. (C) Quantitative comparisons of promoter read coverage (reads per kilobase) of each histone modification using 2.5-kb TSS centered bins. (D) Change in the enrichment of each histone modification bound to a gene promoter plotted against the change in expression of that gene using the set of differentially expressed genes during SSC aging.

#### Figure S5. Genome browser representation of histone modification changes. Examples of individual genes showing the increased (A) or decreased (B) H3K27me3 enrichment
