## Supplementary figures and images for "Transcriptomic and epigenomic profiling of young and aged spermatogonial stem cells reveals molecular targets regulating differentiation"

### Supplemental Figure 1

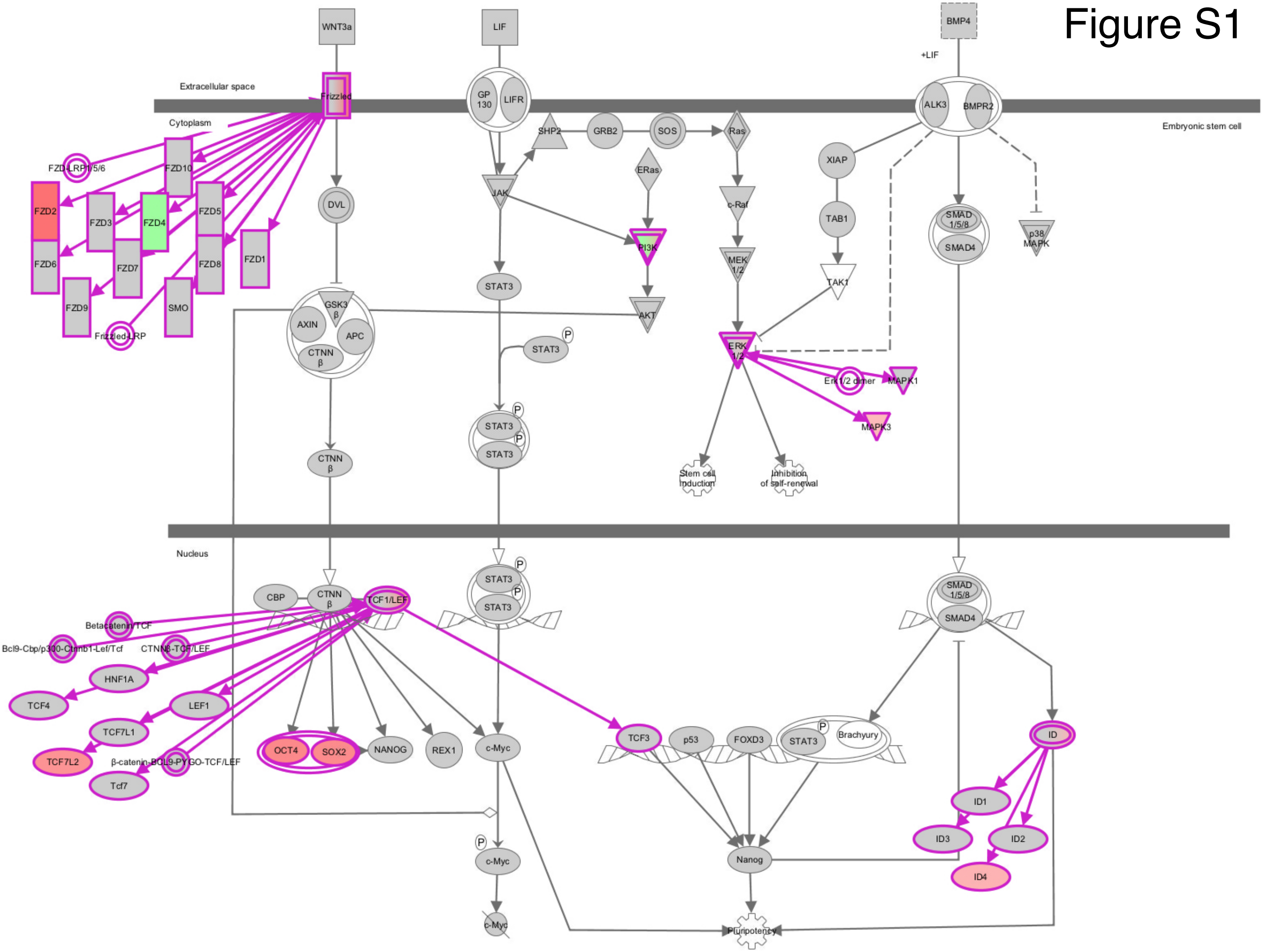

### Supplemental Figure 3

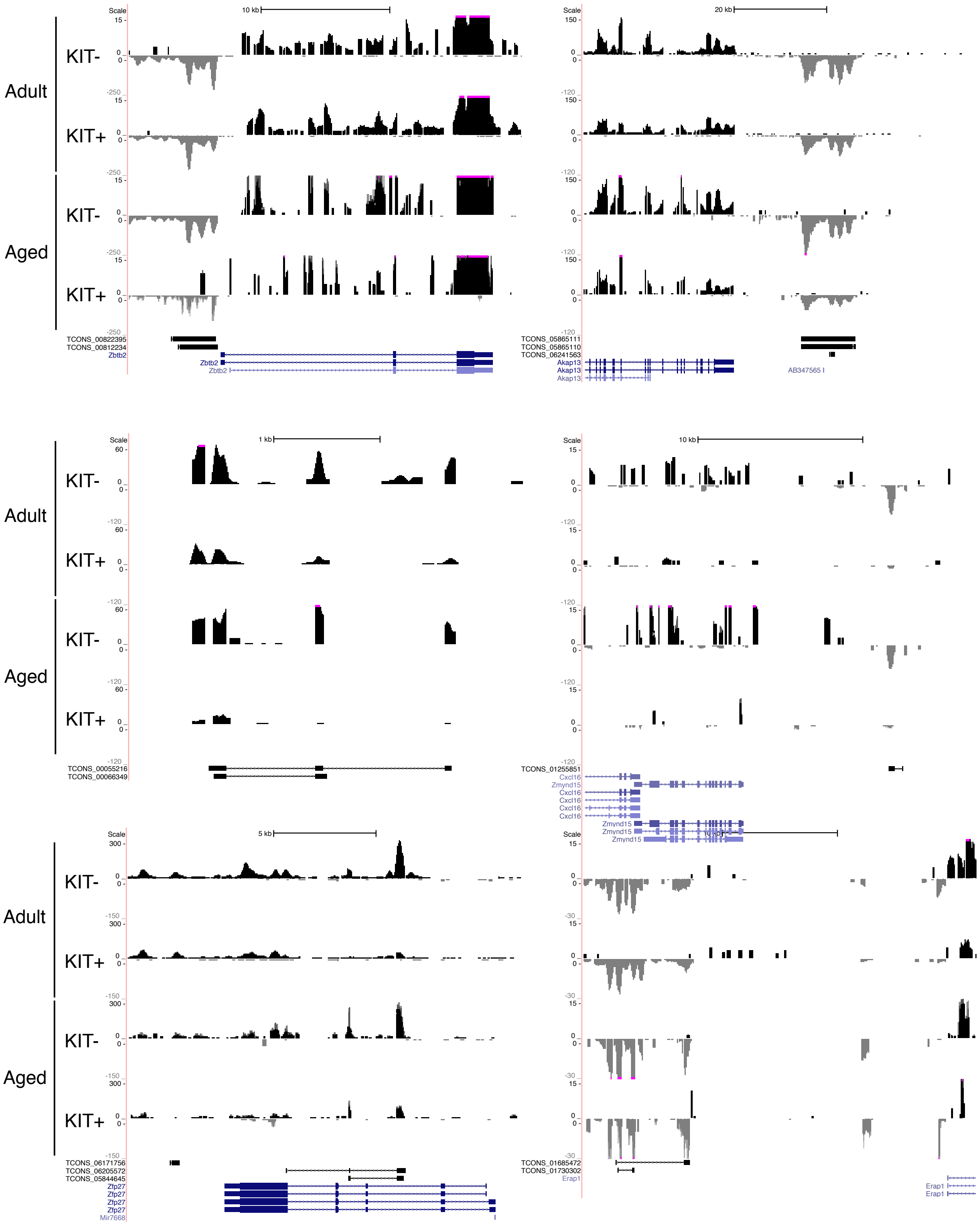

### Supplemental Figure 4

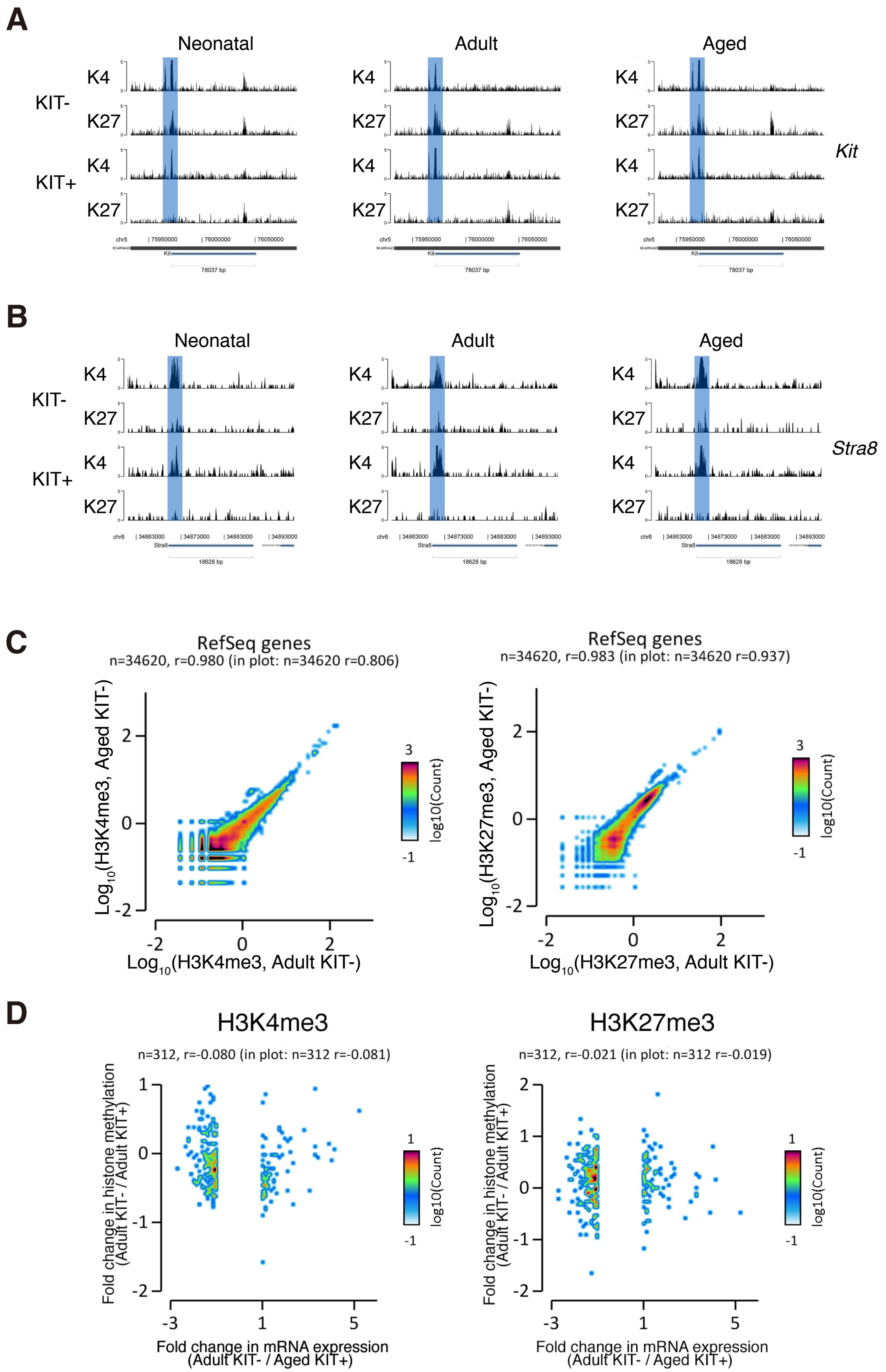

### Supplemental Figure 5

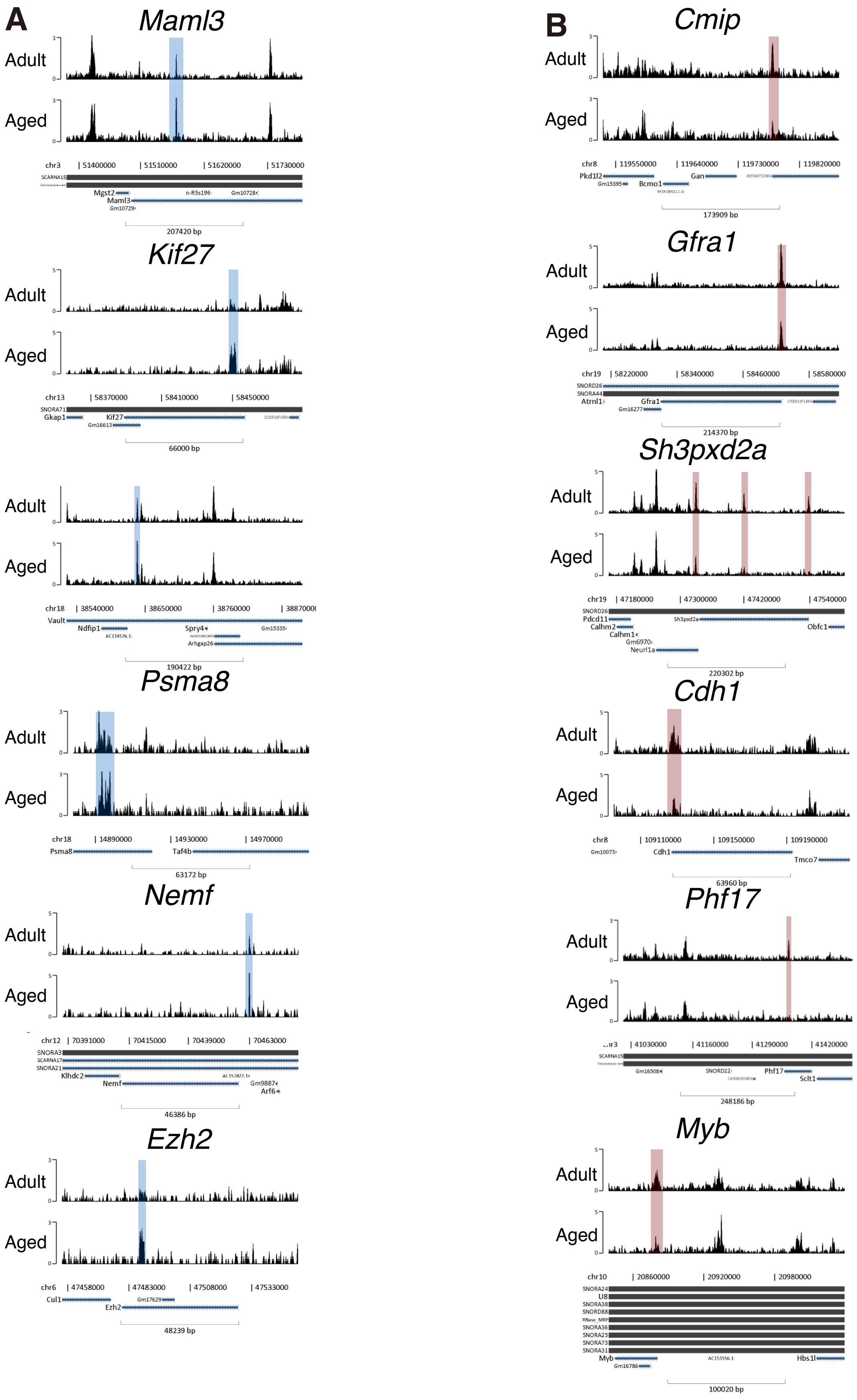
