## Supplemental Figure 2 for "Transcriptomic and epigenomic profiling of young and aged spermatogonial stem cells reveals molecular targets regulating differentiation"

### Figure S2

**A**

| Data Source | This study | GSE49622 | GSE35005 | GSE43717 |
|---|---|---|---|---|
| SSCs age | neonatal | | neonatal | neonatal |
| | adult | adult | | |
| | aged | | | |
| SSCs stage | undifferentiated | undifferentiated | indistinguishable | indistinguishable |
| | differentiating | | | |
| Isolation method(marker) | FACS (Oct4/c-kit) | MACS (Thy1/c-kit) | BSA gradient | BSA gradient |
| Mapped reads | 570 millions | 137 millions | 41 millions | 70 millions |

**B**

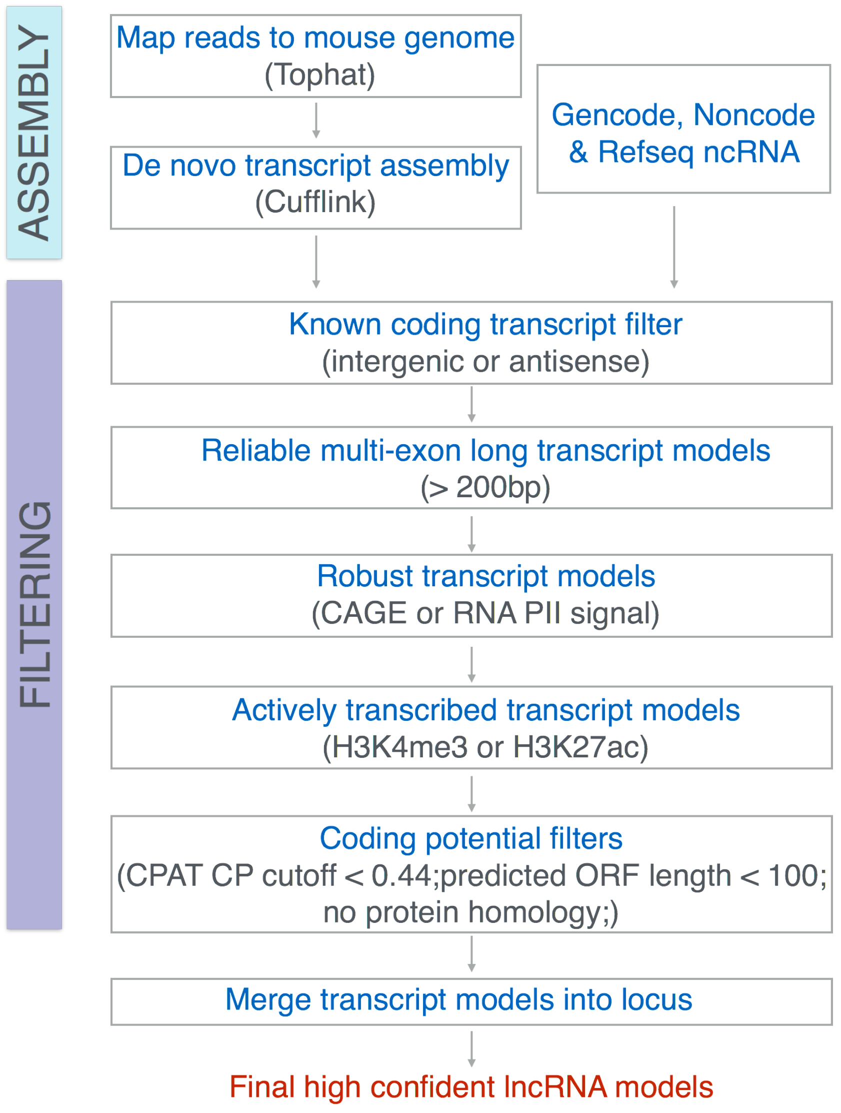
